# Highly efficient, all-organic bioluminescence-photosensitizer conjugate eradicates early-stage tumors and prevents metastasis in mice

**DOI:** 10.1101/2022.01.29.478339

**Authors:** Hao Yan, Sarah Forwad, Kwon-Hyeon Kim, Yue Wu, Jie Hui, Anokhi Kashiparekh, Seok-Hyun Yun

## Abstract

Photodynamic therapy (PDT) is an established treatment modality using light-activatable drugs. Despite its unique cytotoxic mechanism, the shallow penetration of light has been a serious drawback limiting the applications of PDT. Here, we report bioluminescence-activated PDT (BL-PDT) using efficient bioluminescence resonance energy transfer (BRET) conjugates of clinically approved photosensitizers, Ce6, and luciferase proteins. A high photon-to-Ce6 conversion efficiency (80%), along with intracellular delivery by membrane-fusion liposomes, enabled effective cancer cell killing *in vitro*. In a syngeneic mouse model of aggressive 4T1 triple-negative breast cancer, as well as a xenograft model of MDA-MB-231 tumors, BL-PDT resulted in complete tumor remission and prevention of metastasis, as well as neo-adjuvant effects. Our result shows the promise of molecularly activable, clinically viable, depth-unlimited phototherapy.

## Introduction

Photodynamic therapy (PDT) is a clinically approved treatment modality for several diseases including cancers^1-2^. PDT uses light and photosensitizers that, upon absorbing photon energy, can generate reactive oxygen species (ROS) such as single oxygen. The ROS leads to various therapeutic effects, such as the killing of malignant cells, destruction of angiogenic blood vessels, and activation of antitumor immune responses. Effective PDT critically relies on sufficient light dose at target tissues, typically requiring an optical fluence of ∼ 50 J/cm^2^ at the surface of tissue. This amount of dose can produce PDT effects at tissue depths up to a few millimeters below the surface^3-7^. This shallow depth has been an important limitation of PDT, constraining their clinical applications.

To overcome the limited therapeutic depths, a technique to generate light within tissue by Cherenkov radiation has been proposed^8^. However, its use of high-energy radiation and inorganic TiO_2_ nanoparticles raise safety concerns for clinical translation. Another strategy replacing the external light illumination of conventional PDT is using chemiluminescence or bioluminescence (BL) as internal light sources^9-11^. Although the molecular luminescence has much weaker intensity than what external irradiation can provide, efficient Förster resonance energy transfer from the luminescent group to photosensitizers can, in principle, compensate for the intensity difference and produce substantial PDT effects. Recently, a combination of a chemiluminescent donor, luminol, and a photosensitizer as the acceptor was shown to produce efficient ROS-induced cell death in vitro and tumor growth inhibition in vivo^9-10^. However, luminol requires endogenous H_2_O_2_ at a high concentration in the target tissue. This, along with its high affinity to serum albumin and DNA, raises biosafety concerns^11-12^. Compared to chemiluminescence with low quantum yields (< 0.1), bioluminescence using luciferin / luciferase as the donor offers higher quantum yields of > 0.4.^13-14^ Bioluminescence resonance energy transfer (BRET) to Rose Bengal^15^ and photosensitive fusion proteins has been explored^16^, but their low activation efficiencies limit cell-killing effects. BRET using inorganic quantum dots can be more efficient but is not suitable for clinical translation due to the toxic quantum dot materials.

Here we report highly efficient BRET-induced PDT enabled by a novel photosensitizer agent consisting of clinically used photosensitizers Chlorin e6 (Ce6) and Renilla reniformis Luciferase 8 (RLuc8) proteins. An optimized RLuc8 to Ce6 ratio (1:25) and excellent spectral matching between BL emission and Ce6 absorption^9-10^ allowed us to achieve a high activation efficiency — probability of producing an activated Ce6 per BL photon — of 80%. We show that fusogenic nano-liposomes are a highly effective vehicle to deliver the conjugates into cancer cells. We compare the therapeutic effects of our BL-PDT technique with conventional PDT for 4T1 murine and MDA-MB-231 human triple-negative breast cancer (TNBC) cells *in vitro* and then *in vivo* using mouse models. With one-time injection of Luc-Ce6, we demonstrate complete remission of the tumor and prevention of metastasis in lymph nodes and lungs. We also evaluate the neo-adjuvant effect of intratumoral BL-PDT for advanced tumors.

## Results

### Synthesis and BRET efficiency of Luc-Ce6 conjugate

Figure 1a illustrates the working principle of the Luc-Ce6 conjugate. Briefly, we prepared a Ce6 derivative with 6-amino hexanoic acid amide, termed Ce6-HA (710 Da). Ce6-HA molecules are conjugated to RLuc8 proteins (∼37 kDa) using EDC/Sulfo-NHS chemistry (Fig. S1). One RLuc8 protein has a total of 311 amino acid residues^17-18^. The conjugation is formed between the active amino residues and the acid amide of the Ce6-HA, resulting in Ce6 molecules conjugated on the surface of RLuc8. When a substrate Me-eCTZ (briefly CTZ hereinafter) binds with the RLuc8 (briefly Luc8), the BL reaction can occur, producing optical energy, which would otherwise radiate as a photon but in this case, is desired to be transferred to one of the Ce6 molecules via BRET. The excited Ce6 then may react with an oxygen molecule nearby and convert it to singlet oxygen (^1^O_2_). This probability of ROS generation from activated Ce6 is 65-70% in typical *in vitro* and *in vivo* conditions^19-20^.

**Figure 1.**
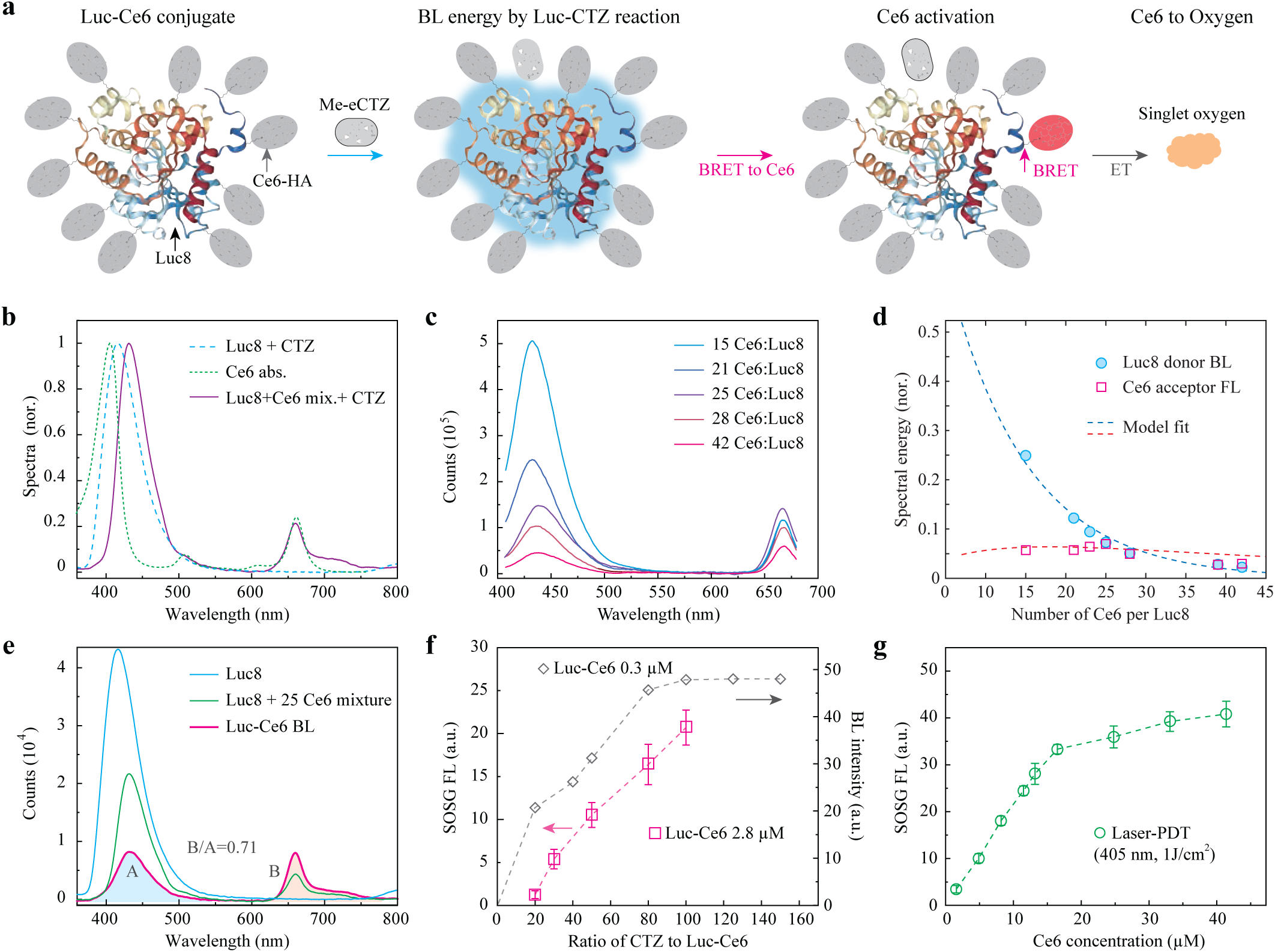
Design and physicochemical characterization of the Luc-Ce6 conjugate. (a) Schematic of the structure of Luc-Ce6 conjugate and its working principle. A Ce6 molecule receives resonant energy transfer from the complex of a RLuc8 protein (Luc) and substrate methoxy e-coelenterazine (Me-eCTZ). (b) Normalized BL spectrum of Luc (blue dashed line), absorption spectra of Ce6-HA (green dashed line), and BL spectra of the mixture of Luc and Ce6-HA (the ratio is 1:25) (purple). (c) BL spectra of Luc-Ce6 conjugates with different mole conjugation ratios of Ce6-HA molecules to Luc protein. (d) The dependence of spectral energy for the donor and acceptor on the number of Ce6 molecules per Luc8. (e) BL spectra of Luc8 protein (0.3 µM), the mixture of Luc (0.3 µM) and Ce6-HA (7.5 µM), and Luc-Ce6 conjugate (0.3 µM for Luc). A: donor emission (400-600 nm) area, B: acceptor emission (600-800 nm) area. (f) Changes in the BL intensity and SOSG fluorescence intensity as the proportion of substrate CTZ is varied at two different concentrations of Luc-Ce6. (g) SOSG fluorescence intensity as a function of Ce6 concentration in solution after laser-PDT (405 nm, 125 mW/cm^2^, 8 s).

A mixture of Luc8 and CTZ in solution emits BL photons with a center at 415 nm, which overlaps well with the absorption peak of Ce6 at 406 nm (Fig. 1b). When Luc8 (0.3 µM) andCe6-HA (7.5 µM) were simply mixed in solution with CTZ, fluorescence from Ce6 at 660 nm appeared (Fig. 1b), due to the absorption of BL photons by Ce6 and BRET from Luc8/CTZ to Ce6 molecules in proximity (< 5 nm) to each other. We fabricated Luc-Ce6 conjugates with different ratios of Ce6 to Luc8. As the ratio increased from 1:1 to 15:1, the BL intensity at 440 nm decreased, and the fluorescence intensity at 660 nm increased. And it had a plateau at ratios between 15 and 28 (Fig. 1c). We determined the optimal conjugation ratio to be 25:1 and used this value in subsequent experiments. A simple kinetic model considering a BRET efficiency and hindrance of Ce6 to the interaction of Luc explained the experimental result reasonably well (dashed lines in Fig. 1d).

Figure 1e shows the representative emission spectrum of Luc-Ce6 conjugates (0.3 µM), integrated over 1 min after injecting a non-saturating small amount of CTZ (∼1 µM). The BL spectra from RLuc8 and a simple mixture of Ce6 and RLu8 with the ratio of 25 are also shown. The BRET ratio, defined by the ratio of total acceptor emission (B, 600-800 nm) to the total donor emission (A, 400-600 nm), was 0.71. Quantitative absorption (Fig. S2b) and fluorescence (Fig. S3b) measurements confirmed the average number of Ce6 in the Luc-Ce6 conjugate to be about 25 as intended. The probability of BL energy ending up exciting a Ce6 molecule can be calculated by using a typical Ce6 fluorescence quantum yield of 15%^21^. The ratio of the number of activated Ce6 molecules to the number of BL photons that are generated without Ce6 conjugation was determined to be 80% (Table S1). This is a remarkably high efficiency of excitation. For far-field illumination, the absorption cross-section of Ce6 is ∼ 1 × 10^−15^ cm^2^ at 405 nm. This means 10^15^ photons / cm^2^ are needed to excite a single Ce6 molecule with a probability of 37% (1/e). In fact, the great majority of photons supplied by far-field illumination in conventional PDT end up not being absorbed by photosensitizers. By contrast, 80% of BL photon is used to excite Ce6.

We used a singlet oxygen sensor green (SOSG) dye^22^ to detect ^1^O_2_ generated by Luc-Ce6 and CTZ in solution. The SOSG fluorescence intensity increased with an increasing amount of CTZ (square, Fig. 1f). In a separate experiment, we measured BL intensity to reach a plateau when the ratio of CTZ to Luc8 exceeded 100 (diamond, Fig. 1f). The result was compared with the SOSG fluorescence after conventional laser-PDT using Ce6-HA as the photosensitizer and 405 nm laser irradiation with an intensity of 125 mW/cm^2^ for 8 s (Fig. 1g and Fig. S5). In terms of singlet oxygen generation, 280 µM CTZ on 2.8 µM Luc-Ce6 is equivalent to laser fluence of 1 J/cm^2^ on 10 µM Ce6.

### Intracellular delivery of Luc-Ce6 conjugates

Intracellular delivery of protein constructs has been challenging^23-25^. We found fusogenic nano-liposomes^24^ (Fuso-lip) to be more effective than conventional liposomal delivery (Fig. S6 and Fig. S7). Upon fusing into the cellular membrane, the liposome can directly inject its cargo into the cytoplasm (Fig. 2a). This direct delivery escapes from the typical endocytosis process in which the protein cargo is confined and rapidly degraded in lysosomes^26-27^. We loaded the Luc-Ce6 conjugate in Fuso-lip using charge differential drive followed by membrane extrusion through 100 nm pores. Using dynamic light scattering, we measured the hydrodynamic size of Luc-Ce6 to be ∼ 17 nm, Fuso-lip to be ∼ 70 nm, and Luc-Ce6/Fuso-lip complex to be ∼ 100 nm (Fig. 2b). The final Luc-Ce6/Fuso-lip complex had a zeta potential of -17.8 mV (Fig. S8).

**Figure 2.**
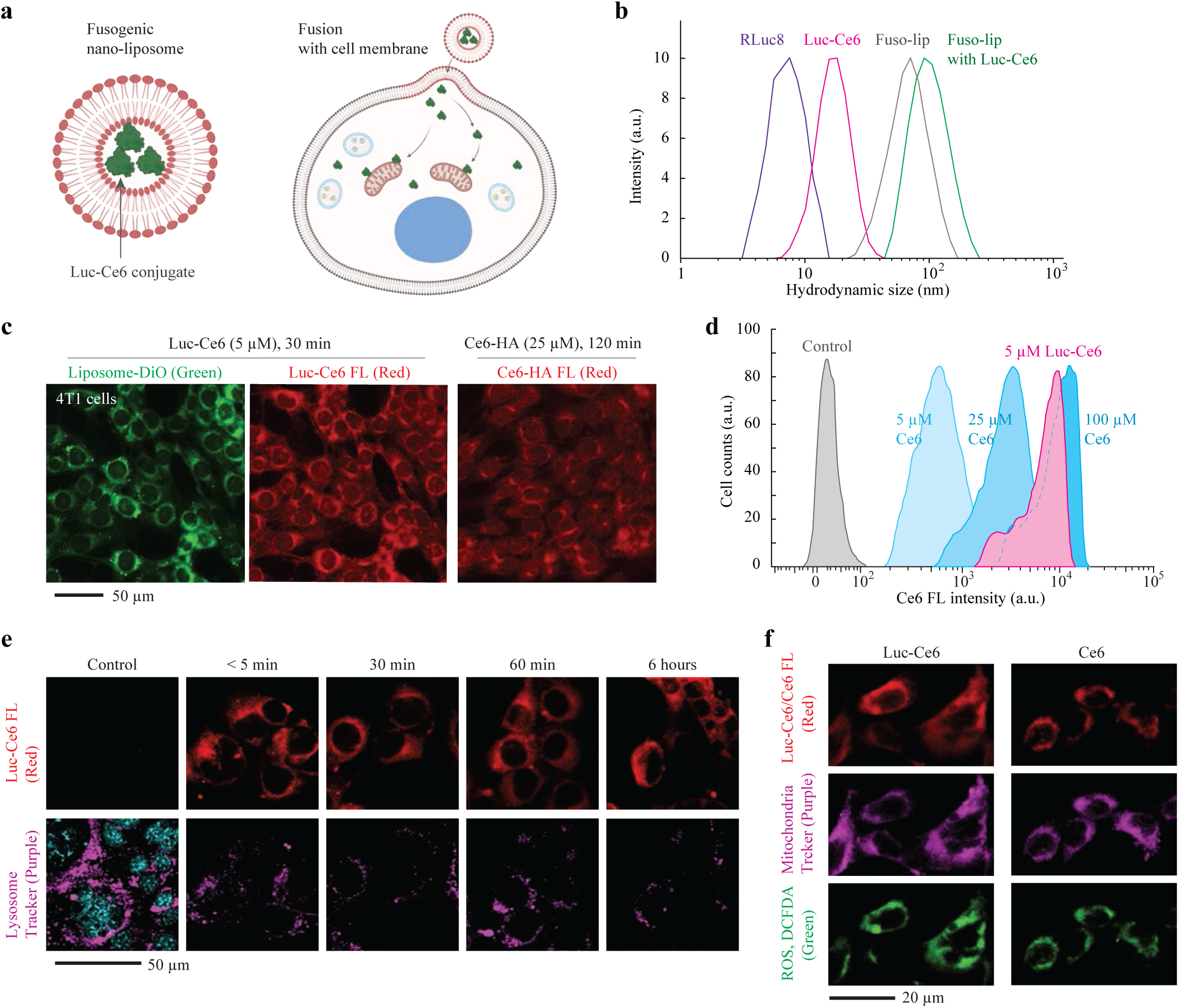
Intracellular delivery of Luc-Ce6 conjugates. (a) Schematic illustration of intracellular delivery. Luc-Ce6 loaded in fusogenic nanoliposomes (Fuso-lip) enters the cell after Fuso-lip is fused with the cell membrane. Luc-Ce6 is in the cytoplasm produces ROS after reacting with CTZ. (b) The hydrodynamic sizes of the Luc protein, Luc-Ce6 conjugate, Fuso-lip, and Luc-Ce6/Fuso-lip. The Luc-Ce6 is about 17 nm in DPBS buffer, and the final Fuso-lip with Luc-Ce6 is about 100 nm. (c) Fluorescence images of 4T1 cells after incubation with 5 µM Luc-Ce6 (red)/Fuso-lip (green) for 30 min, or Ce6-HA molecules (25 µM) for 2 h. (d) Flow cytometry data of cells incubated with 5 µM Luc-Ce6/Fuso-lip and Ce6-HA with different concentrations. (e) Confocal fluorescence images of 4T1 cells after incubation with Luc-Ce6 (red)/Fuso-lip over time. The lysosomes were stained by Lysosome Tracker (purple). (f) Confocal fluorescence images of Luc-Ce6/Fuso-lip-treated and Ce6-treated 4T1 cells. Mitochondria (purple) were stained using a Mitochondria tracker kit, and intracellular ROS (green) was stained using 2’,7’-dichlorofluorescein diacetate (DCFDA).

We tested intracellular delivery with 4T1 murine breast cancer cells. Figure 2c shows fluorescence confocal images of 4T1 cells after 30 min incubation with Luc-Ce6/Fuso-lip (Fig. S9). Fuso-lip fuse immediately with the cell membrane. Luc-Ce6 conjugates are distributed in the cytosol but not in the nucleus. The cells incubated with 5 µM Luc-Ce6 (containing 75 µM Ce6) in Fuso-lip exhibited stronger Ce6 fluorescence than cells incubated with 25 µM Ce6-HA for 120 min incubation time. Flow cytometry analysis supported the effectiveness of Luc-Ce6 delivery by Fuso-lip, nearly comparable to the intracellular uptake of small Ce6-HA molecules (Fig. 2d). The intracellular distribution of Luc-Ce6 was not correlated with the locations of lysosomes (Figs. 2e, S10, and S11). This supports that the uptake mechanism of Luc-Ce6/Fuso-lip escapes from typical endocytosis and does not involve the endosomal-lysosomal system^28-29^. Ce6 molecules are known to accumulate in the mitochondria membrane after entering cells^30-31^ partly because of their hydrophobic aromatic rings. This association with mitochondria is thought to be important for high cytotoxicity of PDT^32^. We observed an overlap of mitochondria with Ce6 and Luc-Ce6 (Figs. 2f, S12, and S13), which also coincide with the fluorescence signal from DCFDA, an intracellular ROS sensor, in both laser-PDT (Fig. S14) and BL-PDT (Fig. S15). Given the similar intracellular distributions, we expect similar cytotoxic mechanisms between BL-PDT and laser-PDT using Ce6.

### *In vitro* BL-PDT on 4T1 and MDA-MB-231 cells

We evaluated the intrinsic cytotoxicity of CTZ, Fuso-lip, and Luc-Ce6/Fuso-lip reagents on 4T1 (murine) and MDA-MB-231 (human) TNBC cell lines and, also, L929 (mouse fibroblast) cells in culture using a Cell Counting Kit-8 (CCK-8) assay (Fig. S16). No significant cytotoxicity was observed from all three cell types for 48 h even with high concentrations of Luc-Ce6 (10 µM) and CTZ (100 µM).

Using the DCFDA intracellular assay, we detected considerable ROS generation in 4T1 cells after BL-PDT (10 µM Luc-Ce6, 100 µM CTZ) (Fig. 3a), which was comparable to conventional PDT (1 J/cm^2^) using 4-times higher Ce6 amount (Fig. 3a). The DCFDA signal increased with increasing amount of Luc-Ce6 for a fixed amount of CTZ (Figs. 3b, S17, and S18). BL-PDT resulted in near complete cell killing confirmed by live/dead fluorescence dyes and morphological changes (Figs. 3c and S19-21). Cell killing by laser-PDT occurs only in regions illuminated with sufficient laser energy (Figs. 3c and S21). When we placed layers of chicken breast tissues on top of cell samples in the optical path, simulating deep tissue therapy, no phototoxicity was observed even with a modest tissue thickness of 300 µm (Fig. S22). Using the CCK-8 assay, we compared the dose dependent cytotoxicity (Figs. 3d and 3e). The half-maximal inhibitory concentration (IC_50_) values of Luc-Ce6 in BL-PDT were measured to be 5.6 µM for 4T1 cells and 1.5 µM for MDA-MB-231 cells. By comparison, IC_50_ values of Ce6 in laser-PDT were 21 µM for 4T1 and 5.4 µM for MDA-MB-231 cells, respectively. Moreover, we used an Annexin V-FITC/PI apoptosis detection kit and flow cytometry to confirm apoptotic death of 4T1 cells 24 hours after BL-PDT and laser-PDT (Fig. S24).

**Figure 3.**
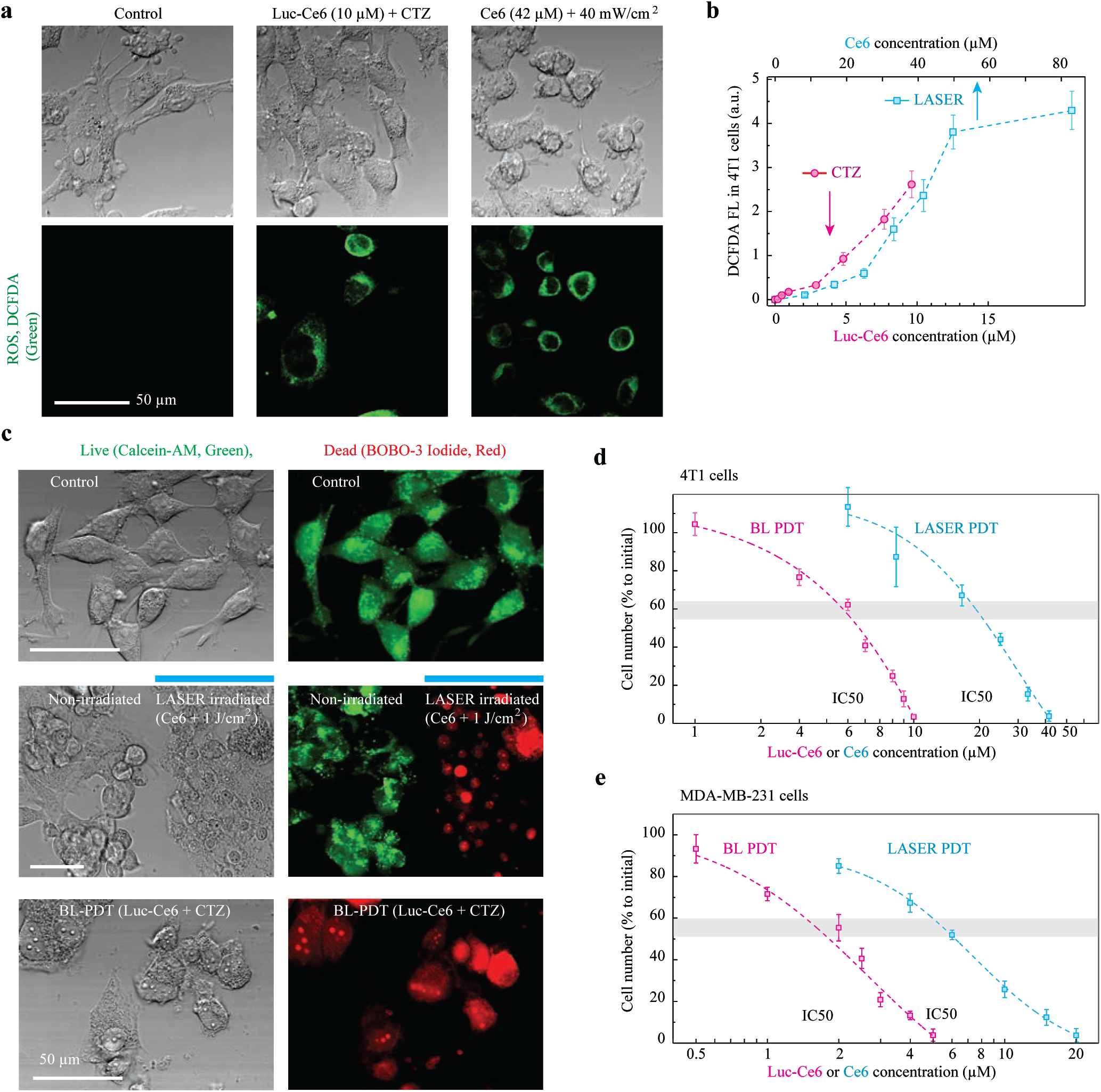
*In vitro* assessment of Luc-Ce6 based BL-PDT. (a) Confocal microscopy images of intracellular ROS-sensing CDFDA (green) in 4T1 cells after BL-PDT (Luc-Ce6 10 µM and CTZ 100 µM) and laser-PDT (Ce6 and 405 nm laser, 1 J/cm^2^). (b) Measured DCFDA intensity for BL-PDT and laser-PDT (1 J/cm^2^) for various concentrations of Luc-Ce6 and Ce6. (c) Fluorescence images of 4T1 cells stained with Calcein AM (live cells, green) and BOBO-3 Iodide (dead cells, red). Scale bars, 50 µM. (d-e) Relative viability of 4T1 (d) and MDA-MB-231 cells (e) after BL-PDT and laser-PDT for different concentrations of Luc-Ce6 and Ce6. The horizontal grey band indicate the half-maximal inhibitory concentration (IC_50_) levels.

### Tumor cell killing by intratumoral BL-PDT *in vivo*

For preclinical testing, we first established a metastasis TNBC orthotopic model, using an injection of ∼10^6^ 4T1 cells into the mammary fat pad of a BALB/c mouse. The 4T1 mammary carcinoma is highly tumorigenic and invasive and, unlike most tumor models, can spontaneously metastasize from the primary tumor to multiple distant sites. The implanted tumor grew rapidly, reaching 4-5 mm in size in 1 week and 9-12 mm in 3 weeks (Fig. S25). Tumor invasion into surrounding tissues was observed at Day 14 (Fig. S25c). Consistently, no apparent metastasis was found in the sentinel lymph node (SLN) and lung at Day 7. On Day 14, small metastases were detected in the SLN and lung (Fig. S26 and Fig. S27). On Day 21, multiple metastatic foci were evident throughout the lung. From this result, we determined that early metastasis occur between 1 and 2 weeks.

We assessed the effects of BL-PDT using a protocol depicted in Fig. 4a. On day 14 after tumor inoculation, Luc-Ce6/Fuso-lip (2.5 mg/kg) was intratumorally injected into the tumor at several sites using a fine needle. To confirm intratumoral distribution, we harvested tumors 6 h after an intratumoral injection of Luc-Ce6/Fuso-lip without CTZ injection and found the reagent distributed extensively in the tumor (Figs. 4b, S28, and S29). For mice in the treatment group, following a Luc-Ce6/Fuso-lip injection we waited for 6 h for the Luc-Ce6 to be delivered to the tumor cells and then intravenously injected the first dose of CTZ (100 µM, 50 µl). Then, 5 additional injections of CTZ with the same dose were given in subsequent 24 h (Fig. 4a). The tumors were harvested at 3 days after the treatment was completed. Their frozen tissue sections were examined using the terminal deoxynucleotidyl transferase dUTP nick-end labeling (TUNEL) assay to detect cell apoptosis. The BL-PDT treated groups showed strong TUNEL signals over larger areas (Fig. 4c). PBS-treated and Luc-Ce6-sham-treated tumors had no obvious TUNEL signals (Figs. 4d and S30). Laser-PDT treated groups, which had received the same intratumoral injection of Luc-Ce6/Fuso-lip and treated with 405-nm laser (90 J/cm^2^) instead of CTZ, showed TUNEL signals only in superficial regions along the tumor boundary within < 1 mm depth from the surface (Fig. 4e). This result showed the advantage of BL-PDT over laser-PDT in terms of therapeutic depth.

**Figure 4.**
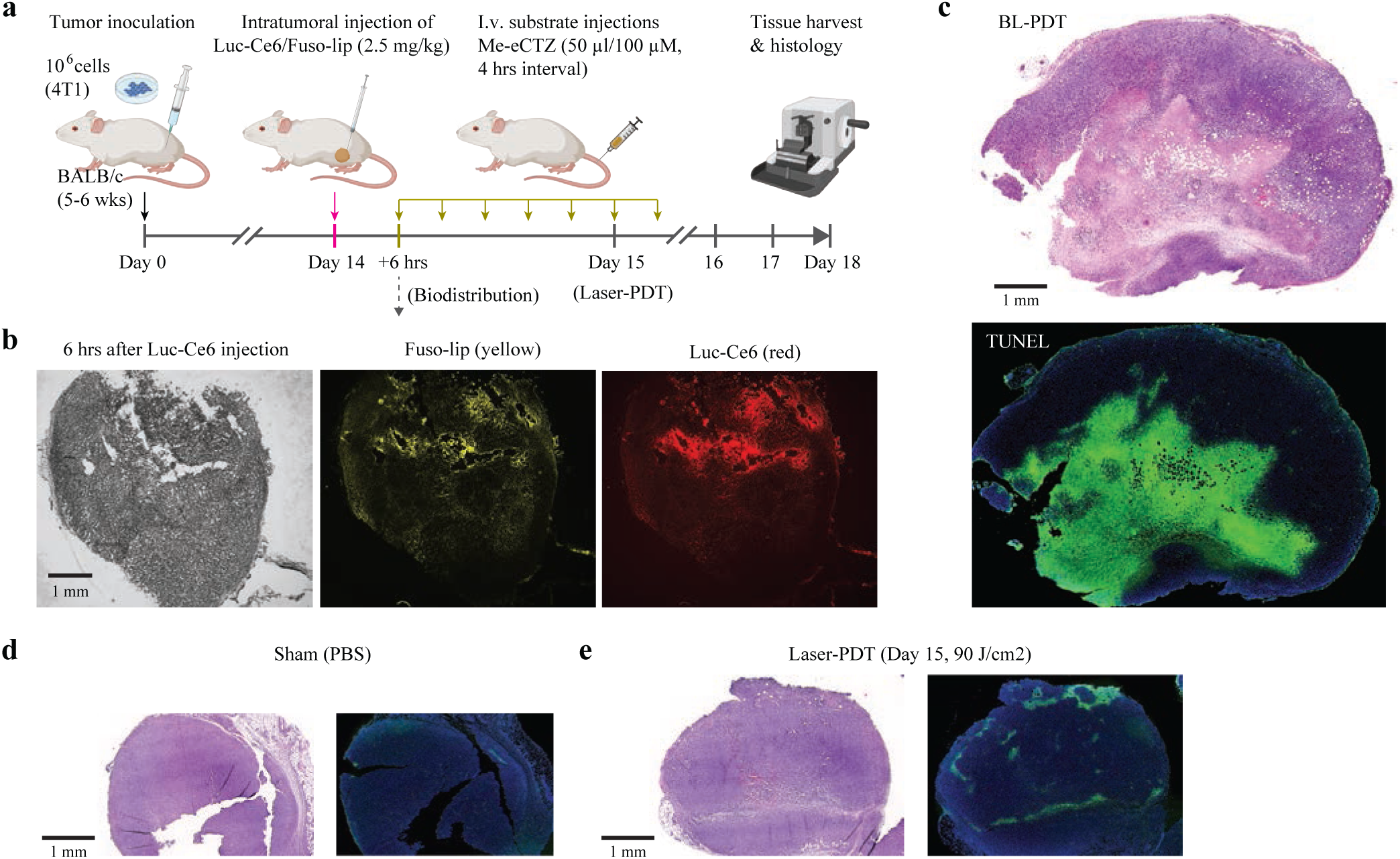
*In vivo* assessment of BL-PDT in a 4T1 TNBC orthotopic model. (a) Schematic representation of the treatment schedule for 4T1 tumors at the advanced stage. (b) Fluorescence microscopy images of the whole tumor cryosection from the Luc-Ce6/Fuso-lip-treated mice. Fuso-lip: yellow, Luc-Ce6: red, BF: bright field. (c) H&E and immunofluorescence images of tumor sections after BL-PDT treatment. Green: TUNEL, blue: DAPI for nuclei. (d) H&E and TUNEL-stained tumor sections after PBS injection. (e) H&E and TUNEL-stained tumor sections after laser-PDT (405 nm, 90 J/cm^2^).

### *In vivo* antitumor treatment by BL-PDT

Having confirmed the cell killing effect *in vivo*, we tested BL-PDT on tumors in their early stage of development. Figure 5a depicts the treatment schedule. BL-PDT was applied on tumors 7 days after 4T1 cell inoculation, a time point when no metastasis is expected to have occurred. For BL-PDT treated mice (n=5), we measured an immediate shrinkage of the tumor volume within 2 days after treatment. A complete tumor regression was achieved by Day 18, without relapse (Fig. 5c). The body weights of the treated mice were normal (Fig. 5d). In stark contrast, laser-PDT treated mice showed nearly unaffected, exponential growth curves (Fig. 5b and 5c). Various organs were harvested at Day 25. Histology of tissue sections from the heart, liver, kidney, and spleen showed no sign of tissue damage (Fig. S31), nor metastasis (Figs. 5f-g and S32). At Day 25, all the control, sham, and laser-PDT treated mice showed substantial metastasis to the SLN and lung (Figs. 5e-g). Our result showed that BL-PDT can eradicate early-stage tumors and prevent metastasis in mice.

**Figure 5.**
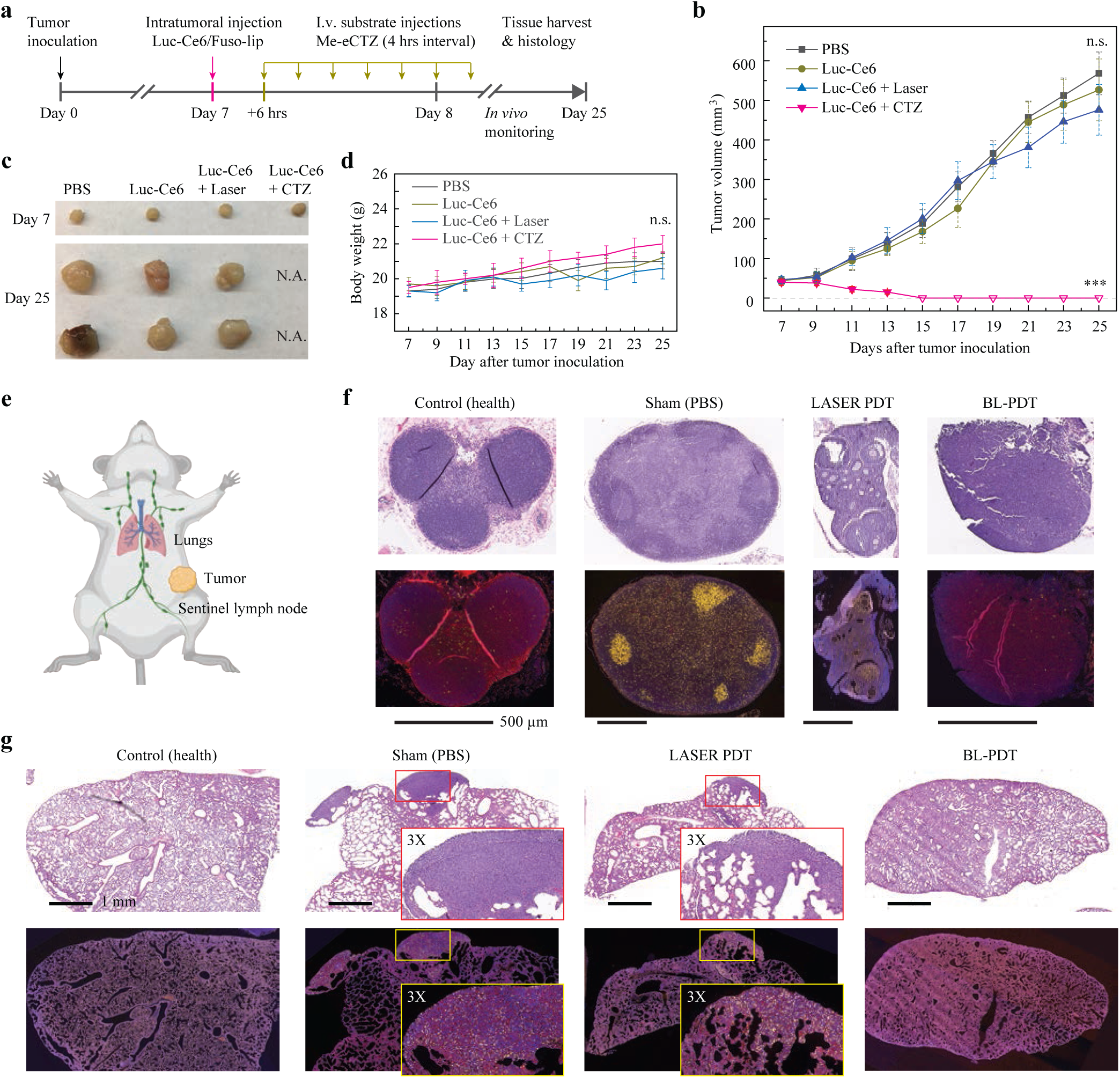
The effects of BL-PDT on 4T1 tumor growth. (a) Treatment schedule. (b) Growth curves of tumors in different groups: PBS injection only, Luc-Ce6 injection only, Luc-Ce6 and laser-PDT (405 nm, 90 J/cm^2^), and BL-PDT. Error bars are the standard deviation of 5 animals per group. ***, P < 0.001; n.s.: not significant. (c) Photos of tumors harvested from different groups at different times. (d) Body weight. (e) Schematic showing the orthotopic tumor and sentinel lymph node. (f-g) H&E histology and immunofluorescence (IF)-stained sentinel lymph nodes (f) and lungs (g) from breast tumor-bearing mice 18 days after treatment: unperturbed control, PBS injection only (sham), laser-PDT and BL-PDT. 4T1 cancer cells were stained by Ki-67 (cancer cell nucleus, yellow), EGFR (cell membrane, red), and DAPI (nucleus, blue). While metastasis is apparent in the lymph nodes and lungs in the PBS and laser-PDT groups, no evident metastasis was found in the control and BL-PDT groups.

### Neoadjuvant BL-PDT

Surgery is the gold standard treatment for removing primary and large tumors. Neoadjuvant chemotherapy and radiation therapy are often performed before surgery to downstage the tumors and reduce the extensiveness of surgery and postoperative complications^33-35^. We tested the neoadjuvant potential^36-38^ of BL-PDT (Fig. 6a). We injected Luc-Ce6/Fuso-lip at the boundary of primary tumor on Day 14, a time point when 4T1 cells typically have started invasion (Fig. 6b). Immediately after the injection, Luc-Ce6 and Fuso-lip are mainly localized in the tumor boundary (Fig. S33). 5 days after the completion of BL-PDT, the tumors were collected for histological analysis. Most tumor cells in the border appeared to be round, atrophic, and apoptotic (Figs. 6c and S34). This morphological change visualized the margin more vividly than tumors in control animals (Fig. 6d) in which the tumor invasion continued to have progressed. Tissues in the laser-PDT treated group also showed blurred tumor boundaries (Fig. S35). Our result shows that adjuvant BL-PDT can shrink the tumor size and delineate the tumor margin.

**Figure 6.**
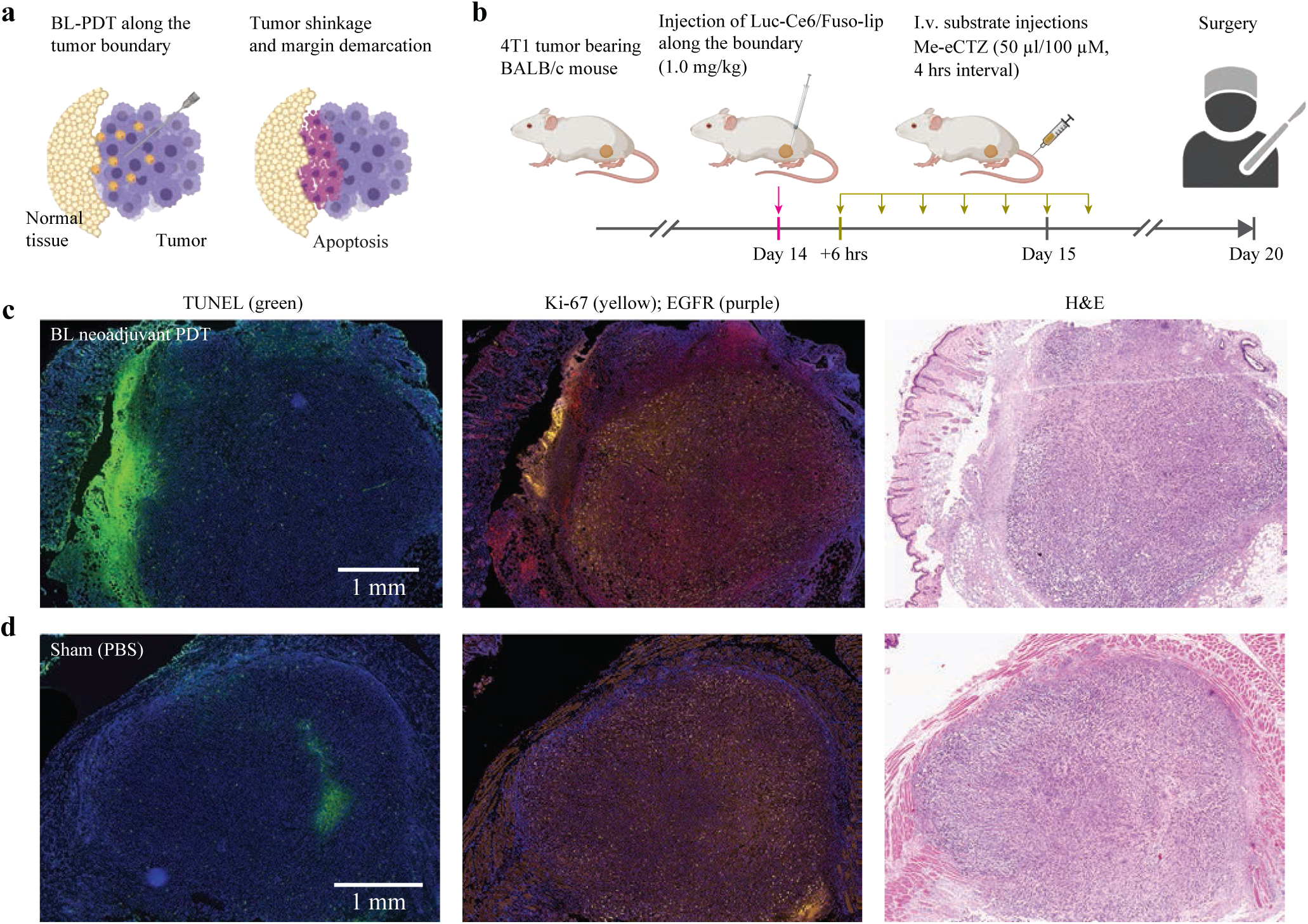
*In vivo* evaluation of BL-PDT for neoadjuvant therapy. (a) Neoadjuvant BL-PDT of an advanced primary tumor. (b) The treatment schedule of neoadjuvant BL-PDT for 4T1 orthotopic tumor model. (c-d) TUNEL, H&E, and immunofluorescence (IF)-stained tumor sections, including surrounding normal tissues, five days after (c) BL-PDT and (d) sham (PBS) treatment. TUNEL: green; Ki-67: yellow; EGFR: purple.

## Discussion

The optimized luciferase-photosensitizer BRET conjugates, along with membrane-fusion liposomal delivery, enabled us to demonstrate highly effective BL-PDT for early-stage treatments of TNBC. The molecularly activatable Luc-Ce6 conjugate enables depth-independent PDT, a major advantage over conventional light-activatable photosensitizers. The toxicity of the all-organic reagent is minimal to none until CTZ is administered. This triggerable action allows BL-PDT to be performed at an optimal time when the agents have been delivered to target regions for the best therapeutic outcome. It could also help achieving low background cytotoxicity, an advantage over non-activable drugs such as conventional chemotherapy agents.

The ROS-mediated mechanism makes PDT a unique modality different from chemotherapies, radiation therapies, and immunotherapies^39-41^. Our results demonstrated the potential of BL-PDT as a standalone, complementary, or synergistic treatment option for cancer treatment. In this study, we have investigated intratumoral delivery by local injection of Luc-Ce6/Fuso-lip. Although this method could be viable for treating small tumors or neo-adjuvant treatment of larger tumors, as demonstrated, it may be a worthwhile to develop a systemic delivery method of BRET conjugates for broader applications such as treating metastases^42-45^. It will also be interesting to develop other combinations of luciferase, luciferin, and photosensitizer than Luc-Ce6-CTZ, with possible differences in target tissues and organelles. BL-PDT can be applicable to treating non-cancers, such as deep skin lesions. Finally, the BRET strategy may be extended to other phototherapies, such as antibacterial blue light therapies^46-47^, that use endogenous photosensitizers. Our promising results encourages further developments toward clinical BRET-based phototherapies.

## Materials and Methods

### Synthesis of Luc-Ce6 conjugate

The Luc-Ce6 conjugate was synthesized by directly conjugating Renilla luciferase 8 (Rluc8, or simply Luc, G-Biosciences) with chlorin e6 mono 6-amino hexanoic acid amide (Ce6-HA) molecules (Santa Cruz Biotechnology) using the standard N-(3-dimethyl aminopropyl)-N-ethyl-carbodiimide hydrochloride (EDC)/N-hydroxysulfosuccinimide (Sulfo-NHS)-activated condensation reaction. Briefly, 1 mg Ce6-HA (1.4 × 10^−6^ mol) powder was thoroughly dispersed in 100 µl dimethyl sulfoxide (DMSO) and then mixed with 400 µl DPBS buffer to make a 0.5 ml, 2 mg/ml solution. Then, 100 µl EDC (2.6 × 10^−5^ mol) and 50 µl Sulfo-NHS (4.6 × 10^−6^ mol) were added into a Ce6-HA solution and reacted for 20 min at room temperature. After this crosslinking reaction, 447 µl 2-mercaptoethanol (55 mM) was added to the solution to quench the excess EDC, and the sample was incubated for 10 min at room temperature. For conjugation, a new modified Ce6-HA solution was added into a Rluc8-DPBS solution, gently pipetted several times, and then reacted for 3 h in an Eppendorf ThermoMixer-C instrument (300 rpm) at room temperature. The ratio of Ce6-HA and Rluc8 was varied to test different mole conjugation ratios. After conjugation, the mixture solution was filtered using Zeba spin desalting columns (7K MWCO) to eliminate unreacted small molecules and obtain the final Luc-Ce6 DPBS solution.

### Characterization of Luc-Ce6 conjugate

An electron-multiplication charge-coupled device (EMCCD)-equipped spectrometer (Newton, Andor) was used to measure bioluminescence (BL) spectra from various solutions. To measure BL spectra from RLuc8 and methoxy e-coelenterazine (Me-eCTZ, or simply CTZ), 1 ml, 0.3 µM CTZ was added to a 1 ml, 0.3 µM Luc-Ce6 DPBS solution in a polystyrene cuvette. The BL emission was collected with 0.1 s exposure and 300 frame averaging. The fluorescence spectra of Ce6-HA solutions (from 1.7 to 24.9 µM) and Luc-Ce6 solution (0.38 µM) were measured by using the EMCCD-coupled spectrometer and a xenon lamp filtered by a monochromator. A microplate reader (Epoch 2, Biotek Instruments) in the spectrophotometer mode was used to measure the absorption spectra of Ce6-HA at different concentrations (from 8.3 to 166 µM) and of Luc-Ce6 conjugates (1.39 and 2.78 µM). A dynamic light scattering instrument (Malvern Zetasizer Nano-ZS, 633 nm) was used to measure the hydrodynamic size and Zeta potential.

### Detection of singlet oxygen in solution

A singlet oxygen sensor green dye (SOSG, Thermo Fisher Scientific), which is highly selective for ^1^O_2_, was used to measure the amount of singlet oxygen in solution. A mixture solution of Luc-Ce6 or Ce6-HA with SOSG (dissolved in methanol) was prepared (10 µM SOSG in the mixture), and CTZ with different molar ratios (20 to 150) was added to BL-PDT sample solutions. The laser-PDT sample solutions were illuminated with 405 nm laser beam at 1 J/cm^2^. The fluorescence spectra of the samples were measured 30 min after treatments.

### Fusogenic nano-liposomes (Fuso-lip) and Luc-Ce6 loading

The two main components of liposomes, 1,2-dioleoyl-3-trimethylammonium-propane (DOTAP) and 1,2-dioleoyl-sn-glycerol-3-phosphoethanolamine (DOPE), were purchased from Avanti Polar Lipids (stored at -80 °C). A 1 mg/ml stock solution of DOPE/DOTAP with a weight ratio of 1:1 was prepared. After evaporating the chloroform solvent, DOPE and DOTAP were resuspended in DPBS buffer at 1 mg/ml concentration. For Luc-Ce6 loading, a 500 µl, 27.8 µM fresh solution of Luc-Ce6 was added into an 800 µl DOPE/DOTAP solution, stirred vigorously for 10 min (300 rpm), and incubated in an ultrasonic bath (Elmasonic, 37 kHz, 30% power) for 5 min at room temperature. To reduce the size of conjugate-loaded liposomes, a mini-extruder instrument (Avanti Polar Lipids) was used to gently extrude the Luc-Ce6/Fuso-lip solution several times through membrane filters with 100 nm pore sizes for *in vitro* experiments or 50 nm pore sizes for *in vivo* experiments. After extrusion, 3,3′-dihexadecyloxacarbo-cyanine perchlorate (DiO) or 1,1′-dioctadecyl-3,3,3′,3′-tetramethyl-lindotricarbocyanine iodide (DiR) was added into the liposome solution with a ratio of 1/15 (w/w) for fluorescent staining of the liposomes. The sample was diluted with DPBS buffer to a desired working concentration.

### Cell culture and *in vitro* cytotoxicity evaluation

Murine fibroblast L929 cell line cells, murine TNBC 4T1 cells, and human TNBC MDA-MB-231 cells were purchased from ATCC (American Type Culture Collection). Cells were cultured in recommended standard conditions. DMEM basic cell media was supplemented with fetal bovine serum (10%, v/v) and penicillin/ streptomycin (1%, v/v), known as “complete media”. Cells were incubated at 37 °C in a humidified atmosphere of 5% CO_2_ and 95% air. For the cytotoxicity studies, cells were seeded on a 96-well plate (∼5,000 cells per well) with 100 µl complete media and cultured for 24 h prior to adding different test reagents: CTZ (25, 50, and 100 µM), Fuso-lip (50, 75, 100, and 150 µg/ml), or Luc-Ce6/Fuso-lip (0.75, 1.25, 1.7, 2.5, 3.3, 5.0, 6.7, and 10 µM). After incubation for 4 h for CTZ and 30 min for Fuso-lip and Luc-Ce6/Fuso-lip, the cells were washed two times by DPBS buffer and cultured in fresh cell media for 24 or 48 h. The cell viability was measured by using a cell counting kit-8 (CCK-8, ApexBio) assay. The absorbance was measured by using a spectrophotometer (Epoch 2, Biotek Instruments).

### Intracellular delivery of Luc-Ce6 conjugate

Cells were plated in glass-bottomed well plates with the same density. After culturing for 24 h, the cell media was removed, a Luc-Ce6/Fuso-lip DPBS solution was added, and the cells were incubated for 30 min. After removing the Luc-Ce6/Fuso-lip solution, the cells were washed three times and cultured in fresh cell media for further experiments or fixed by 4% paraformaldehyde (PFA) for imaging. As a control group, Ce6-HA solution was added to the culture plate and incubated for 2 h before fixing with 4% PFA. The cell nuclei were stained with 4’,6-diamidino-2-phenylindole (DAPI). A lyso-tracker probe (Invitrogen, Red DND-99) was used to stain intracellular lysosomes. 1 mM probe stock solution was diluted to 1 µM in DPBS buffer and added to the cells in glass-bottom plates to reach 50 nM in the media. After 1 h incubation, the cells were washed 3 times with DPBS buffer, fixed with 4% PFA, and examined with fluorescence confocal microscopy. A mitochondria tracker Red CMXRos dye (Invitrogen) and 2’,7’-dichlorodihydrofluorescein diacetate (DCFDA, Cell Biolabs) were used for mitochondrial and intracellular ROS staining, respectively. The Mito-Tracker probe was dissolved in DPBS buffer to prepare 50 nM solution. DCFDA was dissolved in DMSO to form a stock solution (500 µM) and diluted into the Mito-Tracker DPBS buffer to obtain a 5 µM working solution. After incubation with Luc-Ce6/Fuso-lip, 4T1 cells in a glass-bottom plate were washed twice with DPBS and added with the Mito-Tracker /DCFDA-DPBS mixture solution. After further incubation for 30 min at 37 °C, the cells were washed three times, fixed, and stained with DAPI for imaging.

### Measurement of intracellular ROS

4T1 Cells were plated on a glass-bottomed plate. After cultured for 24 h, Luc-Ce6/Fuso-lip or Ce6 DPBS solution was added and incubated for 30 min or 2 h, respectively. After washing twice with DPBS, a 50 nM DCFDA/DPBS labeling solution was added into the cell culture and incubated for 30 min at 37 °C. For the BL-PDT group, CTZ (100 µM) was added, and the cells were incubated for 4 h. For the laser-PDT group, 405 nm laser was illuminated with a fluence of 1 J/cm^2^. After completing the treatment, the cells were washed three times and fixed by 4% PFA for imaging.

### Assessment of therapeutic effects *in vitro*

4T1 (5,000 cells per well) and MDA-MB-231 cells (10,000 cells per well) were seeded in 96-well plates and incubated for 24 h. After washing with DPBS, the cells received Luc-Ce6/Fuso-lip at different concentrations and incubated for 1 h at 37 °C. The cells were washed with DPBS, received 100 µM CTZ, and incubated for 4 h. The cells were then washed and then rested for 24 h at 37 °C. The viability of the cells was measured by using the CCK-8 assay. Control cells underwent the same washing and incubation process without adding Luc-Ce6 and CTZ reagents. For laser-PDT, cells were incubated with a Ce6-HA DPBS solution for 2 h and then irradiated with a 405 nm laser for 1 J/cm^2^. For imaging analysis of cell viability, a Live/Dead™ cell imaging kit (Invitrogen) was used. 4T1 cells were incubated for 24 h, rinsed with DPBS, and stained with the live/dead kit for 15 min at room temperature. The cells were washed with DPBS and fixed by 4% PFA for imaging. For apoptosis assay, 4T1 cells were plated in 24-well plates at a density of 5 × 10^4^ cells per well and cultured for 24 h. After incubation with Luc-Ce6/Fuso-lip for 30 min or Ce6 for 2 h, 4T1 cells were treated with 100 µM CTZ for BL-PDT and 405 nm laser (1 J/cm^2^) for laser-PDT. After incubated for 24 h, the cells were harvested, washed twice with cold DPBS, and then resuspended in DPBS buffer. The cell solution was then mixed and incubated with Annexin-V-FITC and PI (Abcam) working solution for 20 min at room temperature. Flow cytometry (BD FACSAria Cell Sorting System) was used to measure the apoptosis signals from the fluorescent probes.

### Tissue penetration efficiency study *in vitro*

To measure the penetration depth of 405 nm laser, chicken breast tissue slices with a thickness of 50 µm each was prepared through standard freezing tissues and cryosection. Each breast tissue slice was fully spread and put on a glass slide. The glass slides were stacked on top of each other to simulate thicker tissues (50 µm for one slice to 300 µm for 6 stacks). 4T1 cells were prepared in a plate and incubated with Ce6-HA for 2 h. 405 nm laser light with a fluence 1 J/cm^2^ was illuminated onto the first tissue slice on top. The optical attenuation through tissue stacks was measured by using an optical power meter (Thorlabs, PM400). After the laser-PDT, the cells were incubated for 24 h, and their viability was measured by using the CCK-8 assay.

### *In vivo* tumor model

All animal studies and related protocols (2015N000205) have been reviewed and approved by the MGH Institutional Animal Care and Use Committee (IACUC) following the NIH guidelines. BALB/c female mice (5∼6 weeks old) were purchased from Jackson Laboratories. A metastatic TNBC orthotopic model was produced by carefully injecting ∼ 10^6^ 4T1 tumor cells into the mammary fat pad of a mouse. From the Week 1 following inoculation, the tumor volume was measured by external manual examination. The tumor volume was calculated as (length times width^2)/2. Each week post inoculation, tumor tissues, sentinel LNs (SLNs), and lungs were harvested to analyze the extent of tumor invasion and metastasis. Tissue sections were stained with Hematoxylin & Eosin (H&E) and antibodies against Ki-67 and epidermal growth factor receptor (EGFR).

### *In vivo* biodistribution after intratumoral delivery

Mice with orthotopic 4T1 TNBC tumors were randomly divided into two groups (n=3 each). Fuso-lip stained with DiR dyes was used. A Luc-Ce6/Fuso-lip PBS solution was injected into the tumor at several locations to cover the entire volume of the tumor at 2.5 mg/kg dose (based on the Luc-Ce6 amount). Mice were sacrificed at 6 h after injection. The entire tumor was carefully collected and quickly frozen in OCT gel (Tissue-Tek®) for cryosection. To obtain spatial distribution of Luc-Ce6 in the tumor, four tumor slices (8 µm thick each) were cut with an inter-slice distance of 50 µm. The sham-treated control group received the same multipoint intratumoral injection of PBS. The tumor slices were imaged by using a fluorescence microscope (Keyence, BZ-X700).

### *In vivo* antitumor efficacy

For initial assessment, 4T1 tumor bearing mice were randomly divided into 4 groups (n=3 each), each receive PBS, Luc-Ce6/Fuso-lip only, BL-PDT, and laser-PDT. At Day 14 after tumor inoculation, Luc-Ce6/Fuso-lip PBS solution was directly injected into the tumor at multiple locations at 2.5 mg/kg dose. 50 µl PBS was injected for the PBS group. For the BL-PDT group, 50 µl CTZ (100 µM) was intravenously injected into the tail vein at 6 h after the injection of Luc-Ce6/Fuso-lip, and additional CTZ with the same amount was administered every 4 h for total 24 h duration. For the laser-PDT group, the skin directly on top of the tumor was irradiated with 405 nm laser with a dose of 90 J/cm^2^. 3 days after treatment, the mice were euthanized, and the whole tumors were collected and fixed in 4% PFA for 2 days. Then the tumor tissues were transferred for staining for H&E, Ki-67, EGFR, and TUNEL following the standard protocol. The tissue slices were scanned by using a Nanozoomer Slide Scanner (Hamamatsu).

For testing early-stage tumor treatments, mice with orthotopic 4T1 tumors were randomly divided into 4 groups (n = 5 each). At Day 7 after tumor inoculation, Luc-Ce6/Fuso-lip PBS solution was injected into the tumor by 2.5 mg/kg dose. Starting from 6 h after the injection, CTZ (100 µM) was intravenously injected to the BL-PDT group 6 times with an interval of 4 h. A single 405 nm laser irradiation (90 J/cm^2^ at the skin) was given to the laser-PDT group. Following the treatments, the tumor size and body weight, as well as general health conditions, of the mice were measured every other day. 17 days after the completion of the treatment, the mice were euthanized. Sentinel lymph nodes (SLNs) and lungs were collected and fixed with 4% PFA for 1 day and embedded into paraffin for sectioning. Tissue sections of the SLNs and lungs were stained with H&E and antibodies against Ki-67 and EGFR. Additionally, various other organs, such as the heart, liver, kidney, and spleen, were also harvested, fixed by 4% PFA and examined by H&E histology.

### BL-PDT for neoadjuvant therapy

2 weeks post tumor inoculation, mice with orthotopic 4T1 tumors were randomly divided into PBS control, Laser-PDT, and BL-PDT groups (n=3 each). For the laser-PDT and BL-PDT groups, Luc-Ce6/Fuso-lip-PBS solution (1.0 mg/kg) was carefully injected into the tumor boundaries in all sides using a find needle. For distribution measurement, 6 h after the injection, the tumor and its neighboring normal tissues were harvested and quickly frozen in OCT gel for cryo-sectioning. For testing neoadjuvant therapy, 50 µl CTZ (100 µM) was intravenously injected every 4 h 6 times for the BL-PDT group. 90 J/cm^2^ of 405 nm laser was given to the laser-PDT group. 5 days after treatment, tissues including the tumor boundary were harvested and fixed in 4% PFA for staining for H&E, Ki-67, EGFR, and TUNEL.

### Statistical analysis

The data are presented as means ± standard deviation, as noted in each case. The number of samples, n, are provided. Two-way analysis of variance (ANOVA) was used. P < 0.05 was considered to indicate a statistically significant difference. P < 0.05, P < 0.01, and P < 0.001 are indicated with single, double, and triple asterisks, respectively.

## Data availability

The data that support the figures within this paper are available upon reasonable request.

## Acknowledgements

This work was supported by US National Institutes of Health (NIH) grant, 5R01CA192878. We thank Drs. Tayyaba Hasan and Seonghoon Kim for contributions in the initial stage of the project, and Dr. Jenny Zhao and the Wellman Photopathology Core for helping with tissue sectioning, staining, and imaging.

## Author contributions

H.Y. and S.H.Y. designed the experiments. H.Y. and K.H.K. fabricated Luc-Ce6 conjugates. H.Y. and S.F. developed fusion liposomal delivery. H.Y. and A.K. performed biocompatibility assays. Y.W. and J.H. assisted in chemistry. H.Y. conducted in vitro experiments. H.Y. and S.F. established the mouse model. H.Y. performed in vivo experiments. H.Y. and S.H.Y. prepared figures and wrote the manuscript, with inputs from other co-authors.

